# The parallel auditory brainstem response

**DOI:** 10.1101/648659

**Authors:** Melissa J Polonenko, Ross K Maddox

## Abstract

The frequency-specific tone-evoked auditory brainstem response (ABR) is an indispensable tool in both the audiology clinic and research laboratory. Most frequently the toneburst ABR is used to estimate hearing thresholds in infants, toddlers and other patients for whom behavioral testing is not feasible. Therefore, results of the ABR exam form the basis for decisions regarding interventions and hearing habilitation with implications extending far into the child’s future. Currently, responses are elicited by periodic sequences of toneburst stimuli presented serially to one ear at a time, which take a long time to measure multiple frequencies and intensities, and provide incomplete information if the infant wakes up early. Here we describe a new method, the parallel ABR (pABR), which uses randomly timed toneburst stimuli to simultaneously acquire ABR waveforms to 5 frequencies in both ears. Here we describe the pABR and quantify its effectiveness in addressing the greatest drawback of current methods: test duration. We show that in adults with normal hearing the pABR yields high-quality waveforms over a range of intensities, with similar morphology to the standard ABR in a fraction of the recording time. Furthermore, longer latencies and smaller amplitudes for low frequencies at a high intensity evoked by the pABR versus serial ABR suggest that responses may have better place specificity due to the masking provided by the other simultaneous toneburst sequences. Thus, the pABR has substantial potential for facilitating faster accumulation of more diagnostic information that is important for timely identification and treatment of hearing loss.

## INTRODUCTION

The frequency-specific auditory brainstem response (ABR) is an essential diagnostic tool for estimating audiometric thresholds in infants and other patients for whom behavioral thresholds are difficult or impossible to obtain. Accurate threshold estimation is critical to determining the need for auditory prostheses such as hearing aids or cochlear implants, and for enrollment in appropriate habilitation programs. This process needs to occur quickly because earlier intervention promotes better spoken speech and language outcomes in children (Ching et al., 2014; Cullington et al., 2017; Harrison, Gordon, & Mount, 2005; Joint Committee on Infant Hearing, 2007; May-Mederake, 2012; Moeller, 2000; Yoshinaga-Itano, Sedey, Coulter, & Mehl, 1998). The toneburst ABR has been the gold-standard for infant assessment because testing can be completed while the infant sleeps and its thresholds highly correlate (up to 0.9) with behavioral thresholds when a large range of thresholds is considered (Gorga et al., 2006; Ramos, Almeida, & Lewis, 2013; Stapells & Oates, 1997). While effective at estimating hearing thresholds, the diagnostic ABR suffers from an important constraint: test time. We aim to describe and validate the feasibility of our new parallel ABR (pABR) method which is designed to address this time constraint by presenting multiple frequencies in both ears simultaneously.

Reducing test time is important for two main reasons. A diagnostic ABR exam entails measuring a series of individual responses at several frequencies over a range of intensities in both ears (American Academy of Audiology, 2012; Hood, 1998, p. 98; Hyde, 2008). Because the exam is highly sensitive to movement artifacts, the ABR is typically performed while the infant sleeps. This constrains the duration of the test to that of the infant’s nap, which also makes the endpoint unpredictable. To compensate, audiologists must make decisions about which frequencies and intensities are the most important to acquire in which ears and pursue those first (e.g., BC Early Hearing Program, 2012, p. 18). If the infant wakes up earlier than anticipated, the audiologist is forced to choose between inferring thresholds from incomplete data or scheduling a return visit in which the test can be completed. This delays diagnosis and treatment, poses risks for attrition, carries additional costs, and adds stress to the family as they await clinical decisions. This is not a trivial burden of time: approximately 150,000 infants are referred for the exam each year in the United States alone, with about 10,000 found to be deaf or hard-of-hearing (Task Force on Newborn and Infant Hearing, 1999; Vohr, 2003). Reducing the exam time and the need for additional visits will free up clinician time and resources, lowering the barrier for referral and increasing the likelihood that children receive the needed care in a timely manner. Accurate threshold estimates must be obtained—early intervention in patients with elevated thresholds leads to improved language, cognitive, and educational outcomes later in childhood.

The second reason test times need to be shortened is to minimize exposure to sedation and anesthesia. While newborns are able to sleep during the exam, infants over four months old and young children often cannot sit still or sleep, requiring the use of sedation or general anesthesia (François, Teissier, Barthod, & Nasra, 2012; Hood, 1998, p. 122). Recent studies investigating the effects of anesthesia on the developing brain suggest a risk of neurotoxicity. Even a few hours of exposure can result in significant neuronal loss in young animals, with deleterious effects persisting later in life (Wagner, Ryu, Smith, & Mintz, 2014; Jevtovic-Todorovic et al., 2003; Creeley et al., 2013; Brambrink et al., 2012). In children, the risk of learning disabilities increases with longer accumulated exposure to some drugs (Wilder et al., 2009). Based on these findings, the FDA issued a warning that general anesthesia and sedation drug use should be avoided or minimized wherever possible for children under three years of age, and should be limited to three cumulative hours (FDA, 2017). Diagnostic ABRs routinely last between 1 to 3 hours, using up the recommended exposure times for the first three years of life. Shortening the diagnostic ABR exam would reduce dosages, and in some situations may make it possible to run the test without drugs. Thus, the imperative to reduce test time extends beyond cost and convenience: it is essential for reducing the risk of damaging the developing brain.

The signal-to-noise ratio (SNR) of an ABR measurement principally drives how long the measurement takes because an ABR waveform is the averaged responses to several thousand repetitions of a stimulus. The SNR of the averaged waveform improves as the number of stimulus repetitions increases. Thus, an attractive way to attempt shortening test time involves increasing the rate at which stimuli are presented. Increasing the stimulus rate in practice, however, carries important drawbacks. With typical periodic stimulus presentation, the time window in which the ABR waveform can be viewed is limited to the inter-stimulus period, or the inverse of the stimulus rate. Relevant ABR components can have latencies of 12–15 ms, which practically limits the rate to 70–80 Hz. Several studies have sidestepped this constraint by replacing periodic stimulus timing with various types of jitter or randomization schemes, allowing stimulation rates into the high hundreds of hertz or beyond (Eysholdt & Schreiner, 1982; Özdamar & Bohórquez, 2006; Valderrama et al., 2012, 2014; Wang, Zhan, Yan, Bohórquez, & Özdamar, 2013). These high rates reduce noise, but neural adaptation shrinks responses (i.e., signal), which in turn partially or completely cancels out SNR gains that the faster rates might have provided. Timing randomization shows some benefit at rates in the low hundreds of stimuli per second, but a lack of translation to the clinic suggests those are outweighed by the increased complexity of analysis and a lack of normative data. Additionally, these studies have focused on clicks, rather than more diagnostically relevant toneburst stimuli.

Recording time could also be reduced if responses to different frequency bands could be recorded simultaneously. This is the approach taken by the multiple auditory steady-state response (ASSR). A single ASSR stimulus is constructed by modulating a sinusoid carrier at the audiometric test frequency. Rather than the waveform, the response is a single automatically derived number (e.g., an F-test) that quantifies the neural phase-locking to the modulator. Giving each test frequency an independent modulation rate allows separate assessment of responses to several simultaneously presented stimuli. The ASSR can effectively reduce test time and estimate thresholds that correlate with behavioral thresholds (Luts, Desloovere, & Wouters, 2006), but it also carries drawbacks. The ASSR does not provide the response waveforms that audiologists are experts in interpreting. The way the stimuli are constructed also leads to much higher energy than equivalent toneburst stimuli, which means the clinician must be careful not to expose the patient to potentially dangerous levels. Despite its availability in a number of clinical devices, the ASSR has seen narrower adoption than the frequency-specific ABR as a diagnostic exam in the clinic.

The goal of this paper is to provide proof of principle for a new paradigm for measuring the ABR to all frequencies in both ears in parallel. The parallel pABR is accomplished through designing stimuli comprised of simultaneous, independently randomized sequences of toneburst stimuli. First, we validate that the paradigm yields high quality canonical brainstem responses at stimulus levels ranging from high to very low, suggesting the pABR’s utility for estimating audiometric thresholds. These responses exhibit standard ABR morphology, minimizing the need for clinician retraining. We then show that the time to reach a satisfactory SNR and residual noise value is better for parallel presentation than for the same randomized toneburst trains presented in serial, especially at lower intensities. Taken together, these findings demonstrate the pABR’s feasibility to meaningfully reduce diagnostic test time with few drawbacks.

## METHODS

### Human subjects

Experiments were completed using a protocol approved by the University of Rochester Research Subjects Review Board (#66988). All subjects gave informed consent prior to participation and were compensated for their time. We collected data from 10 subjects (5 females) with a mean ± SD age of 22.6 ± 4.6 years (range: 18.3 to 34.2 years old). Pure-tone audiometric screening confirmed normal hearing thresholds (≤ 20 dB HL) for each subject at octave frequencies between 500 and 8000 Hz. Subjects self-reported no other neurological abnormalities.

### Stimulus construction

Figure 1 depicts stimulus construction for the pABR. As an overview (details given in the next sections), pABR stimuli are constructed from windowed tonebursts centered at octave frequencies from 500 Hz to 8000 Hz. For each frequency, a toneburst train is created by placing tonebursts randomly within a 1 s epoch. This is repeated for all other frequencies with independent random processes controlling the timing. All toneburst trains are summed, and the process repeated with new random processes for the other ear, comprising a stimulus epoch. Because of the independent timing, we can separately compute the average ABR waveform to each toneburst train from the same electroencephalography (EEG) data free from interference. The stimulus presentation rate and intensity can be varied, and we can compare pABR acquisition to single-frequency serial acquisition by presenting all or only one of the toneburst trains, respectively, from a stimulus epoch.

**Figure 1.**
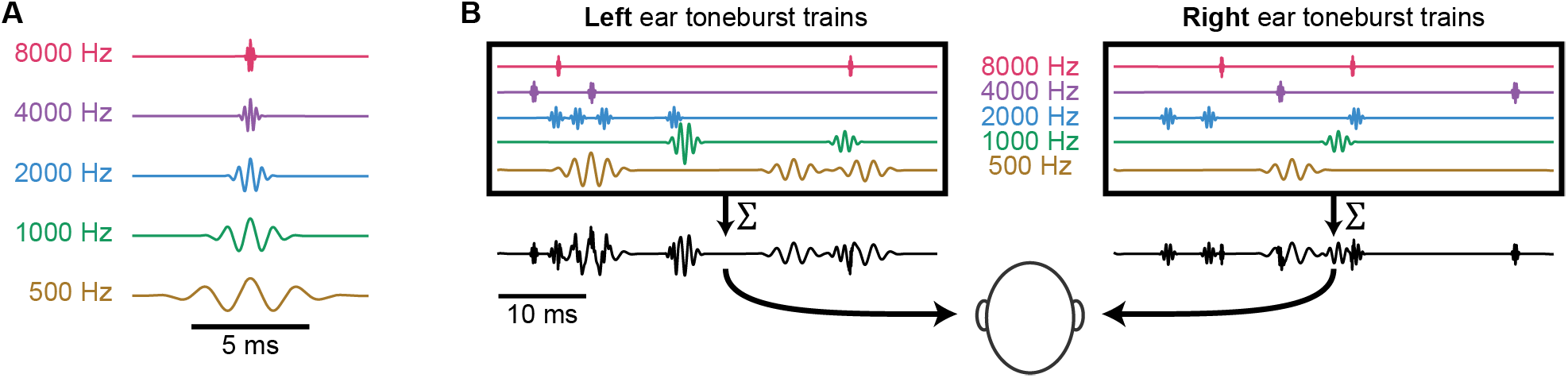
pABR stimulus construction. (**A**) Individual toneburst stimuli for each frequency. (**B**) Toneburst trains in each ear (colored lines) are summed to create a two-channel (left, right) stimulus epoch (black lines).

#### Toneburst stimuli

Toneburst stimuli were constructed at frequencies of 500, 1000, 2000, 4000, and 8000 Hz. For each frequency, five cycles of a cosine were multiplied by a Blackman window of the same length, such that the peak of the window was aligned with a maximum of the cosine function (Figure 1A). Consequently, individual tonebursts had durations of 10, 5, 2.5, 1.25, and 0.625 ms for each of the five frequencies, respectively. Stimuli were generated at a sampling rate of 48 kHz to ensure that the highest stimulus frequencies were well below the Nyquist rate. Stimuli were represented in memory as 32-bit integers so that the full dynamic range could be tested without risk of quantization distortion.

A 1000 Hz sinusoid test tone was used to calibrate the amplitude of the toneburst stimuli. The tone was played from the tube of the insert earphones (ER-2, Etymotic Research) into the sound level meter (2240, Bruel & Kjaer) using a 2cc acoustic coupler (RA0038, G.R.A.S.) and its digital amplitude was adjusted so that its intensity read 80 dB SPL. The amplitude of this sinusoid served as the reference for matching amplitudes of the toneburst cosine components to give a toneburst stimulus level of 80 dB peak-equivalent SPL (peSPL). Other stimulus levels (L) in dB peSPL were obtained by multiplying the reference-level toneburst by 10^(*L* – 80)/20^.

#### Toneburst trains with randomized stimulus timing

Toneburst trains for each frequency were formed by creating an impulse train with random timing at an overall rate of 40 stimuli / s, and then convolving the impulse train with the toneburst. To construct an impulse train, a vector of zeros was first created with a length of 48,000 samples, corresponding to a 1 s interval. Of these, 40 unique sample indices were chosen at random and the zero replaced randomly with +1 or −1 so that half of the tonebursts in each train were inverted. The impulse train was then convolved with the toneburst, creating a toneburst train with half condensation tonebursts and half rarefaction. Indices of the impulse train too close to the end of the 1 s interval were excluded as possibilities if the toneburst would be truncated. This way each epoch had 40 stimuli but no tonebursts were cut off by the end of the epoch.

This process of generating each impulse train timing sequences was essentially a one-dimensional homogeneous Poisson point process, with only very subtle differences. Those differences were: 1) the number of stimuli was set exactly to 40, rather than setting the process’s rate parameter (typically denoted as *λ*) to 40; 2) the indices were guaranteed to be unique (though they could have been at adjacent samples, or 21 μs apart); 3) because the epochs were 1 s long, the maximum inter-stimulus interval was < 1 s. Figure 2 compares the actual inter-stimulus interval histogram of all toneburst trains with the theoretical exponential distribution of an ideal Poisson process with *λ* = 40 stimuli / s, demonstrating that they are practically identical.

**Figure 2.**
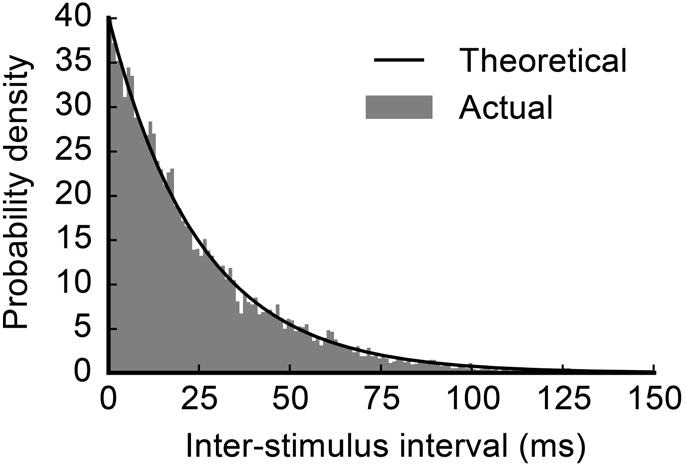
The distribution of inter-stimulus intervals over all stimuli for *λ* = 40 stimuli / s (solid gray) compared to the predicted distribution given by *P*(*t*) = 40*e*^−40*t*^. There is a very close match indicating that the deviations from a true Poisson point process used in this experiment are negligible.

#### Stimulus epochs

Stimulus epochs lasting 1 s were composed of a combination of 10 toneburst trains (5 frequencies × 2 ears). All toneburst trains for the left ear were summed to create the left channel of the stimulus epoch, and the same was done for the right ear (Figure 1B). Each toneburst train was created with a different random seed, such that the timing between any two sequences was completely independent. This statistical independence is what underlies the ability to present stimuli in parallel while acquiring separate responses for each ear-frequency combination.

Thirty unique stimulus epochs were generated to ensure sufficient statistical independence between the random processes dictating the toneburst trains (i.e., the impulse trains that were convolved with the tonebursts) for all frequency-ear combinations. Perfect independence between random sequences is achieved with infinite durations. However, modeling undertaken before data were collected determined that 30 s sequence durations are enough that any channel interactions are far overpowered by the noise endemic to EEG recording. We used a frozen set sequences for which statistical independence was confirmed.

#### Stimulus artifact mitigation

During construction of the stimulation sequence, we employed a double counter-phasing scheme. First, as described above, the polarity of a random half of the tonebursts in each train were inverted, akin to alternating polarity in periodic stimulation. Second, each of the 30 stimulus epochs were followed in the stimulation sequence by an inverted version of that epoch. Thus, the order of the first six stimulus epochs in the sequence was A^+^, A^−^, B^+^, B^−^, C^+^, C^−^, etc., where A, B, and C denote independent stimulus epochs and the superscript denotes the phase.

We also took physical measures to prevent stimulus artifact. We hung earphones from the ceiling so that they were as far from the EEG cap as possible. We also used active cancellation, wherein each earphone attached to another in the same orientation, but with a blocked tube. The dummy earphone received an inverted signal, in order to cancel electromagnetic fields everywhere but close to the transducers. In our experiments this method outperformed passive shielding in artifact reduction, but we note that we have made high-quality recordings without using the dummy earphone method. We also point out that this scheme can be employed in the laboratory, but clinics likely will not (and need not) adopt it.

#### Interleaving trial order

To avoid biases introduced by slow changes in recording quality (e.g., due to changes in subject state or drifting electrode impedances) we interleaved the conditions and consecutively stepped through the trial order. This prevented issues like transient periods of higher EEG noise or slow impedance drifts from singularly affecting one condition over the others.

### Stimulus presentation and EEG recording

Scalp potentials were recorded with passive Ag/AgCl electrodes. A positive (non-inverting) electrode was placed just anterior to the vertex at FCz in the standard 10-20 coordinates and plugged into a y-connector which was split into two differential preamplifiers (Brainvision LLC, Greenboro, SC). The two reference (inverting) electrodes were placed on the left and right earlobes (A1 and A2 respectively). The ground electrode was placed at Fpz. Data were recorded at a sampling rate of 10 kHz and high-pass filtered at 0.1 Hz during recording, with additional filtering occurring from 30 to 2000 Hz offline using a causal first order Butterworth filter.

Subjects sat in a comfortable recliner in a darkened sound-treated room (IAC, North Aurora, IL, USA). They were encouraged to relax and to sleep—nearly all subjects slept for at least part of the test, though this was not rigorously measured. All stimuli were presented through insert earphones (ER-2, Etymotic Research, Elk Grove, IL) which were connected to a stimulus presentation system consisting of a sound card (Babyface, RME, Haimhausen, Germany) and a headphone amplifier (HB7, Tucker Davis Technologies, Alachua, FL, USA). A python script controlled stimulus presentation using publicly available software (available at https://github.com/LABSN/expyfun). Digital triggers were sent from the stimulus presentation computer to BrainVision’s PyCorder software using the sound card’s digital audio out connected to a custom trigger box (modified from a design by the National Acoustic Laboratories, Sydney, NSW, Australia) to precisely mark the start of each stimulus epoch.

### Stimulus conditions used in this study

Stimulus level and presentation rate both have important effects on brainstem responses. These factors and their interactions, as well as optimal ranges, are well studied for traditional ABR. However, the effects of simultaneous stimulation across all frequencies with random timing are not obvious. For this proof-of-concept paper we characterized how the responses to pABR stimulation change over an intensity range, and how these responses compare to those from serial presentation at both a high and low intensity.

In one session we measured responses in both ears to pABR stimulation with an average presentation rate of 40 stimuli / s and intensities in 10 dB steps between 75 and 25 dB peSPL, for frequencies between 500 and 8000 Hz. For a single recording session of 114 minutes, this afforded 16 minutes of recording time per intensity (96 minutes total) to collect 10 responses (5 frequencies each in 2 ears). Three minutes of clicks were also recorded at each intensity but were not analyzed here. Consequently, each averaged response comprised 38,400 repetitions.

In a second session we again measured responses to pABR stimulation at a presentation rate of 40 stimuli / s. We also recorded responses at interleaved trials to a serial single-frequency condition that used the same toneburst trains but tested each frequency separately. We recorded the pABR and serial ABR to frequencies between 500 and 8000 Hz at both a high and low intensity (75 and 45 dB peSPL). To make this possible in a single session, only the right ear was tested in the serial condition, under the rough but necessary assumption that the left ear would show the same behavior. For a single recording session of 108 minutes, this afforded 35 minutes of recording time to collect 20 responses with the pABR (5 frequencies in 2 ears at 2 intensities; 15 minutes at 75 dB peSPL and 20 minutes at 45 dB peSPL for a total of 36,000 and 48,000 repetitions respectively), and 69 minutes to collect 10 responses serially (5 frequencies in 1 ear at 2 intensities; 26 minutes at 75 dB peSPL and 43 minutes at 45 dB peSPL). More time was allocated for recording serially collected responses to low frequency tonebursts and the lower intensity stimuli, such that recording time per serial condition ranged from 4 minutes (9,600 repetitions) for high intensity and high frequency stimuli (i.e., 2000, 4000, 8000 Hz at 75 dB peSPL) to 15 minutes (36,000 repetitions) for 500 Hz at 45 dB peSPL. Four minutes of clicks were also recorded at each intensity but were not analyzed here.

Of the 10 total subjects, 2 were able to complete only one of the two sessions, resulting in 9 subjects for each experiment. For the first experiment, one subject’s recording was too noisy to see responses and so was excluded, resulting in a final total of 8 subjects for the first session and 9 subjects for the second session.

### Data analysis

#### Response calculation

During recording, triggers marked the beginning of each 1 s stimulus block, rather than sending a trigger for each toneburst stimulus. This was for two reasons: 1) random stimulation at overall high rates (when all channels are added together) would have resulted in trigger overlaps, and 2) blocks of stimuli can be analyzed in the frequency domain, which makes calculations substantially faster.

Raw EEG data were bandpass filtered between 30 and 2000 Hz (causal, first order Butterworth filter), and then notch-filtered at odd multiples of 60 Hz in that range to remove power line noise. For each stimulus epoch we calculated a single average response to the 40 toneburst stimuli. Rather than calculate the average response directly, however, we used the mathematically equivalent method of cross-correlation, implemented in frequency domain, between the stimulus sequence and the EEG data. Figure 3 demonstrates this process. Due to the random nature of the stimuli, we were able to extend the analysis window for each toneburst to be 1 s long, which is much longer than the tens of milliseconds duration for the standard ABR (limited to the reciprocal of the presentation rate). This extended window allowed us to calculate response waveforms for the time period 500 ms before and after each stimulus (i.e., from −500 to 500 s, where *t* = 0 is the time of stimulus/toneburst onset). To do this, for each 1 s stimulus block we took the corresponding period of EEG data (1 s) along with the data 500 ms before and after it, leading to 2 seconds of EEG data, denoted as *y*. Then, for each toneburst train, we created a timing sequence by placing a single-sample unit-height impulse corresponding to the start of each toneburst (i.e., from the rectified impulse train created during stimulus construction), and zero-padded it with 500 ms before and after, leading to a 2-second impulse train with all of its impulses located in the middle 1 s, denoted as *x_f,e_*, where *f* is the toneburst frequency and *e* is the ear stimulated. The response waveform, *w_f,e_*, was computed as the circular cross-correlation of *x_f,e_* and *y*, done in the frequency domain for efficiency, as

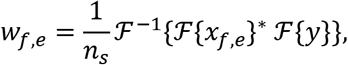

where 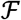 denotes the fast Fourier transform, 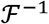 its inverse, * denotes complex conjugation, and *n_s_* the number of impulses in the sequence. The result is a response where the time interval [0, 500] ms is found at the beginning of *w*_*f,e*_ and the interval [−500, 0) ms is found at the end, such that concatenating the two (end first) yields the response over the interval [−500, 500] ms (the middle 1 s of *w*_*f,e*_ is discarded). This process was repeated for each of the ten toneburst trains (5 frequencies, 2 ears) for each epoch. This equation assumes *y* is a single EEG channel, a common scenario for ABR, but this analysis can simply be repeated for each channel if more than one is present. In this study we recorded from two channels, but then averaged the calculated responses for further analysis because we were not concerned with ipsilateral versus contralateral differences for the purposes of this paper. However, separately analyzing ipsilateral and contralateral responses for clinical applications would be easy to perform by keeping the two channels separate rather than averaging. It should also be noted that the typical per-stimulus epoching and averaging in the time domain could have been employed and yields identical results but at greater computational cost.

**Figure 3.**
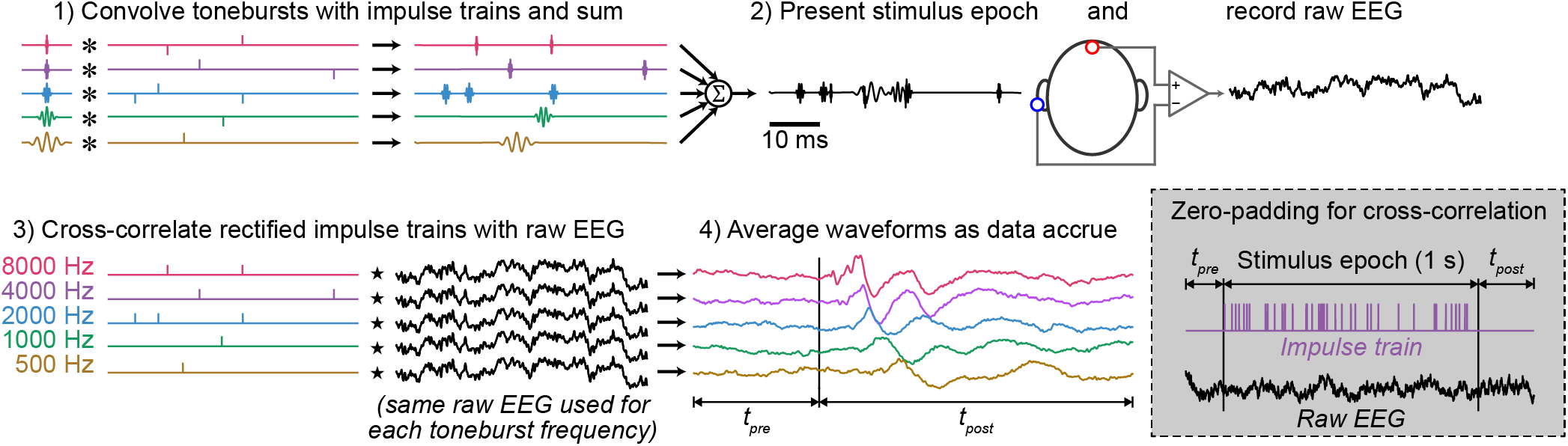
The analysis chain, shown from stimulus creation and presentation to calculation of response waveforms. For clarity, only a 50 ms time period is shown. Dashed box: Zero-padding scheme shown for a single impulse train of a single epoch. Note *t_pre_* and *t_post_* are not shown to scale.

#### Response averaging

Because the quality of the ABR waveforms as a function of acquisition time was of interest, we calculated the cumulative averaged response after each 1 s stimulus block. To account for variations in noise levels over time (either slow drifts or due to transient sources like movement artifacts), we weighed each response according to the inverse of the noise in that epoch. This process is the same in principle as Bayesian averaging described by Elberling and Wahlgreen (1985), but the noise variance was calculated differently. We computed the variance of the pre-stimulus window in the time period −480 ms to −20 ms. We then weighed each epoch by the inverse of its variance relative to the sum of the inverse of variances of all epochs

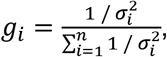

where *i* is the epoch number and *n* is the number of collected epochs. The averaged response was then calculated as

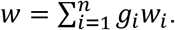

This averaging process avoids the need for artifact rejection based on thresholds, and also takes advantage of the long pre-stimulus window afforded by randomized timing sequences to give a better estimate of the noise.

#### SNR calculation

The SNR of a waveform was estimated by comparing the variance (i.e., mean-subtracted energy) of the waveform in the 10 ms latency range starting at a lag that captured wave V for that frequency (500 Hz: 10.5 ms, 1000 Hz: 7.5 ms, 2000 Hz: 6.5 ms, 4000 and 8000 Hz: 5 ms; Stapells, 2011). That period contained signal and noise, so its variance is denoted 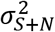. We estimated the noise variance, 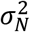, by segmenting the pre-stimulus baseline between −480 and −20 ms into 10 ms intervals, finding the variance of each one, and computing the mean. We then computed the SNR in decibels for every waveform as

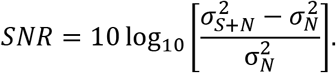

## RESULTS

### The pABR yields canonical waveforms that characteristically change over a range of intensities

We recorded the pABR over a range of stimulus levels from 75 to 25 dB peSPL in 10 dB steps. Figure 4 shows the grand average and responses from two example subjects. Overall response morphology strongly resembled those yielded by traditional methods.

**Figure 4.**
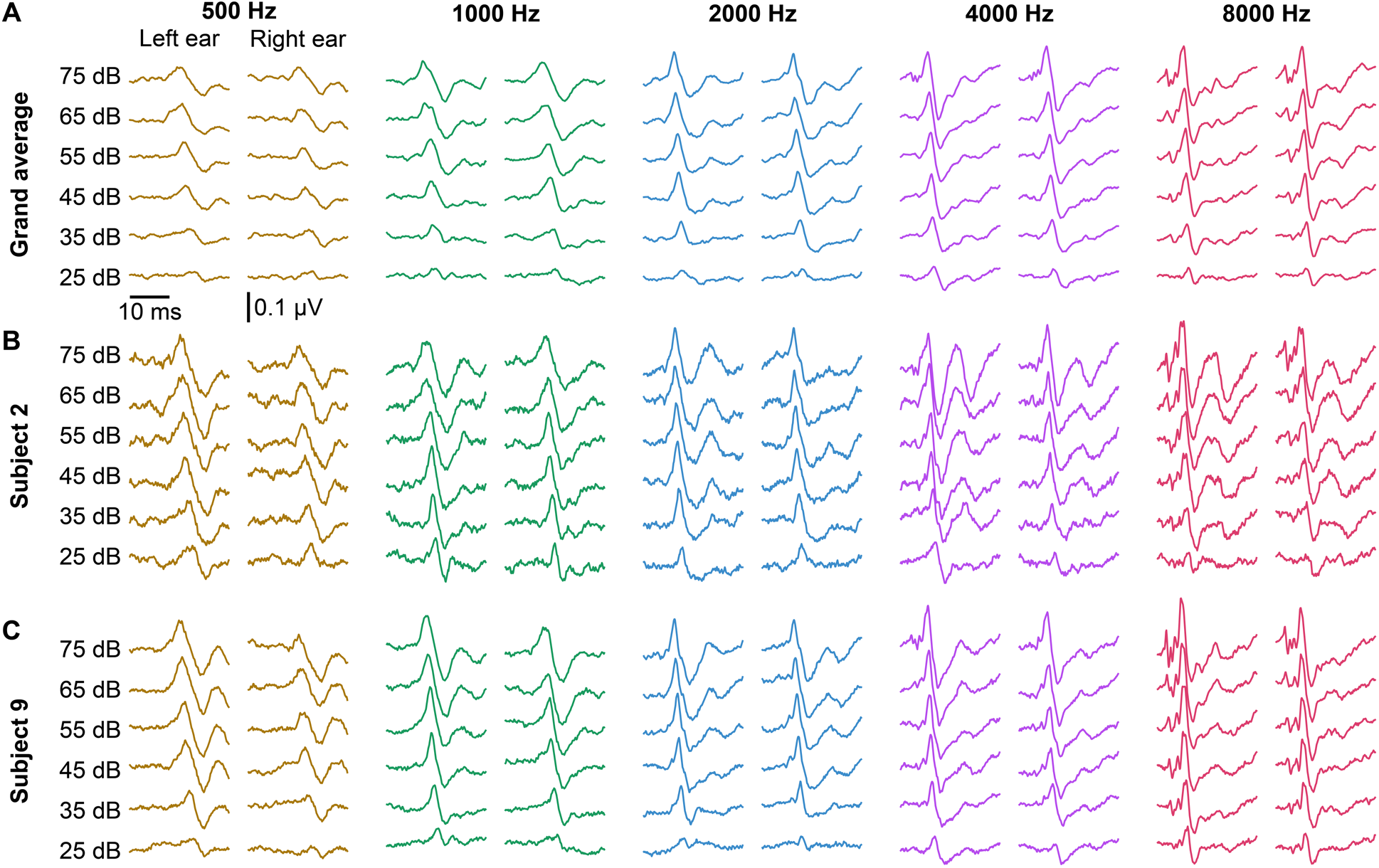
Intensity series waveforms across frequencies and for the left and right ears. (**A**) Grand average of 8 subjects. (**B,C**) Two example subjects’ responses. All responses are plotted over the interval 0 to 25 ms.

Aspects of response morphology were quantified by a trained audiologist (MJP) who manually inspected each waveform to determine the presence, amplitude and latency of wave V. The same measures were quantified by the other author (RKM) in 38% of responses. The intraclass correlation coefficient (ICC3) for each frequency and measure was ≥ 0.9 (all *p* < 0.001), indicating excellent reliability for chosen wave V peak latencies and amplitudes. We modeled wave V latency (Figure 5A) and amplitude (Figure 5B) using two linear mixed effects models, each with a random intercept for each subject and fixed factors of ear, stimulus level, stimulus frequency in log units, and the interaction for log frequency and stimulus level. Wave V latency showed no difference between ears (*p* = 0.66) but there were significant effects of level, frequency, and a significant level-frequency interaction (all *p* < .001), indicating that latency decreased with increasing level and increasing frequency, and the effect of intensity was greater at lower frequencies. Wave V amplitude increased with stimulus level (*p* < .001) and this increase was greater for higher frequencies (significant level-frequency interaction, *p* < .001). These trends are clearly visible in Figure 5, and are generally consistent with traditional ABR (Burkard, Don, & Eggermont, 2006).

**Figure 5.**
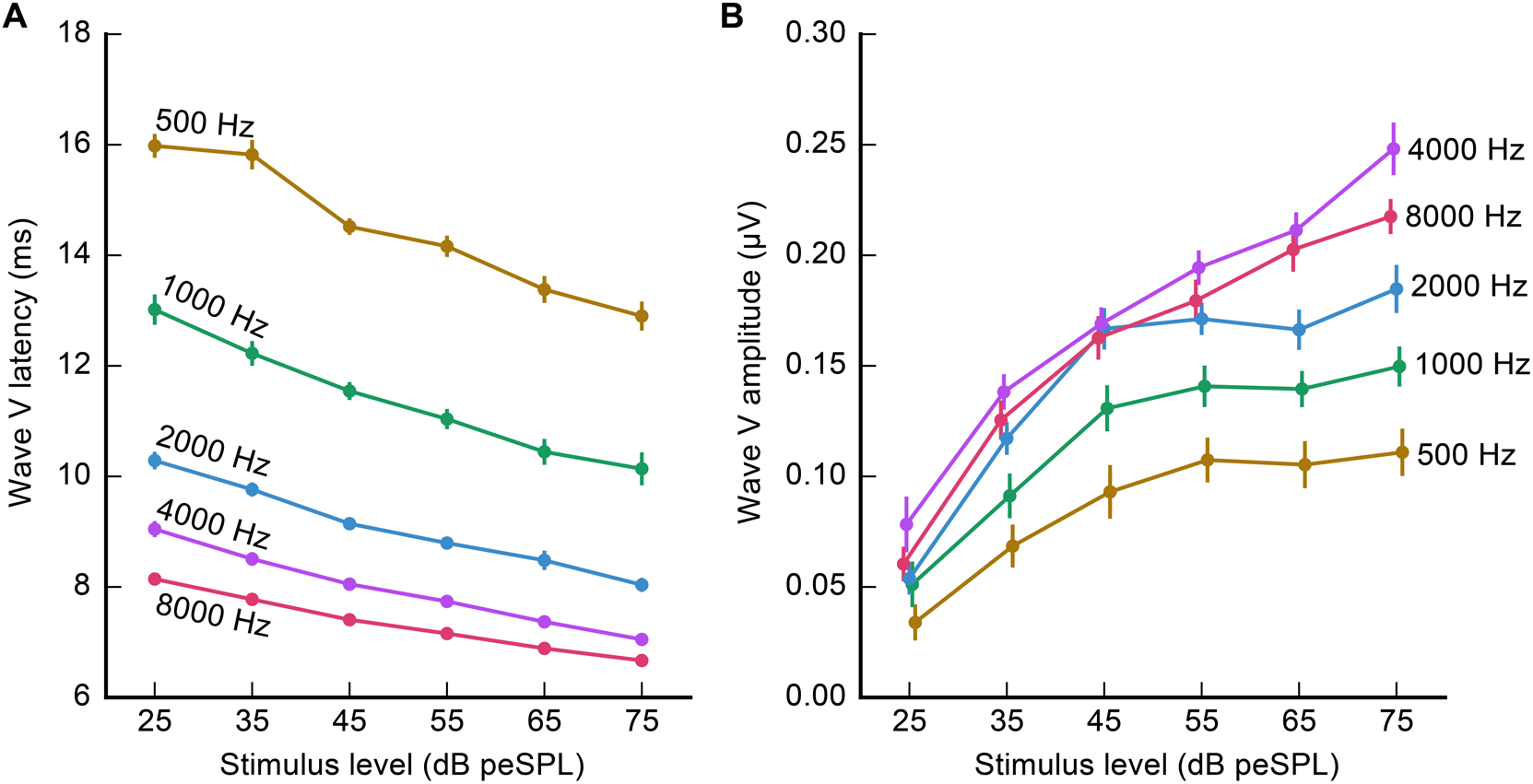
Mean wave V latency (**A**) and amplitude (**B**) as a function of intensity. Stimulus frequency is indicated on each line. Error bars (where large enough to be seen) indicate ±1 SEM. Lines have a slight horizontal offset in **B** to reduce overlap.

While we did not quantify the presence of waves other than wave V, Figure 4 also shows that both waves I and III are clearly visible at higher frequencies in the grand average as well as typical individual responses. Measuring wave I rapidly and at the moderate intensities used in this study may have applications to the study of hidden hearing loss (Liberman, Epstein, Cleveland, Wang, & Maison, 2016).

We thus found that the pABR gives typical ABR waveforms over a range of frequencies and intensities and recapitulates the effects of stimulus frequency and intensity on response morphology seen in traditional ABR. In the next section we directly compare pABR with serially recorded responses recorded in the same subject in a single session.

### pABR and serial response waveforms differ in latency and amplitude at high intensities

We recorded responses in 9 subjects (8 of whom also participated in the previous experiment) to stimulus trains presented in parallel (all frequencies, both ears), versus the same stimulus trains presented serially (one frequency, one ear). Due to time constraints, serial responses could only be recorded in one ear (right), and so even though the pABR recorded responses in both ears, only the right ear responses were compared. Responses were measured for a high and low intensity (75 and 45 dB peSPL respectively).

Figure 6 shows grand averaged responses and responses from two subjects for pABR (colored as in other figures) and the corresponding serial responses (black). Each overlapping waveform is a response to the same stimuli that only differ in the presentation context (parallel, with other stimuli simultaneously present, versus serial, with stimulus trains presented in isolation). Overall waveform morphology of responses were similar using both methods, with some differences in wave V amplitude and latency, described in detail below.

**Figure 6.**
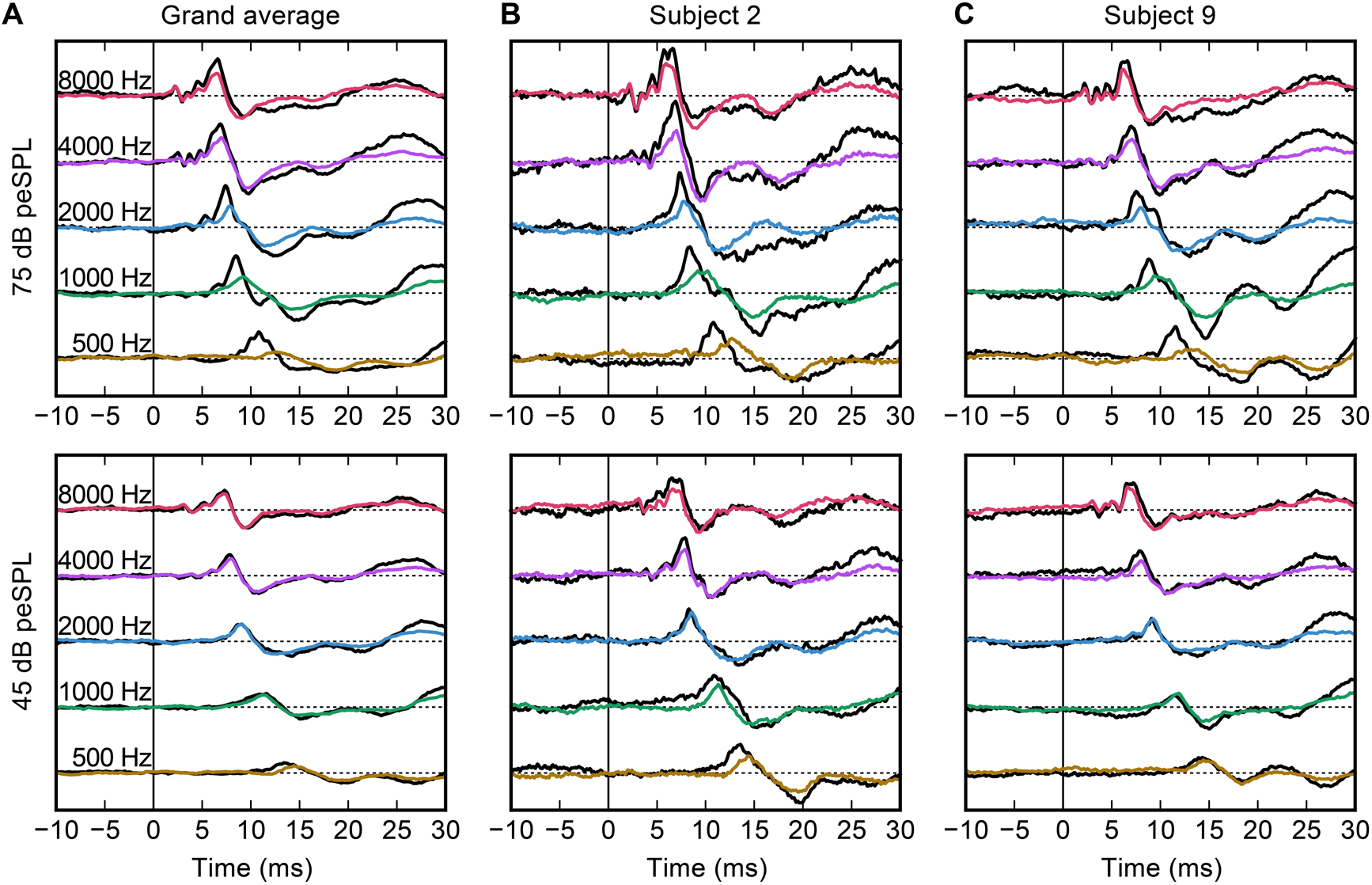
Parallel vs series acquisition waveforms (right ear only). (**A**) Grand average. (**B,C**) Example subjects. pABR is shown in colored lines. Corresponding serial waveforms are shown in black. Vertical spacing is 0.3 μV/ div.

Wave V peak latency and amplitude were further quantified and are displayed for pABR versus serial ABR acquisition in Figure 7. Again, we showed good agreement in our wave V choices (all ICC3 ≥ 0.89, *p* < 0.001). Linear mixed effects models of wave V latency and amplitude were used again with a random intercept for subject and fixed factors of method (pABR versus serial), stimulus level (75 and 45 dB peSPL), log frequency, as well as the full set of two and three-factor interactions. Latency (Figure 7A) showed significant effects of stimulus level (*p* = .012) and frequency (*p* < .001) as well as the method-level-frequency interaction (*p* = .004). Thus, as expected, latencies were longer at lower frequencies and levels. In addition, latencies were longer for pABR than serial ABR for lower frequencies at higher levels. This interaction trend is also clearly visible in the 500 Hz 75 dB peSPL waveforms of Figure 6 for the grand averages and both example subjects. For amplitude (Figure 7B), only the two-way interaction of method and level was significant (*p* = 0.032), indicating that serial ABR shows larger wave V amplitudes at the higher stimulus level. The significant interaction terms of the latency and amplitude models are both consistent with potentially improved place specificity afforded by pABR, a notion which receives a fuller explanation in the Discussion section.

**Figure 7.**
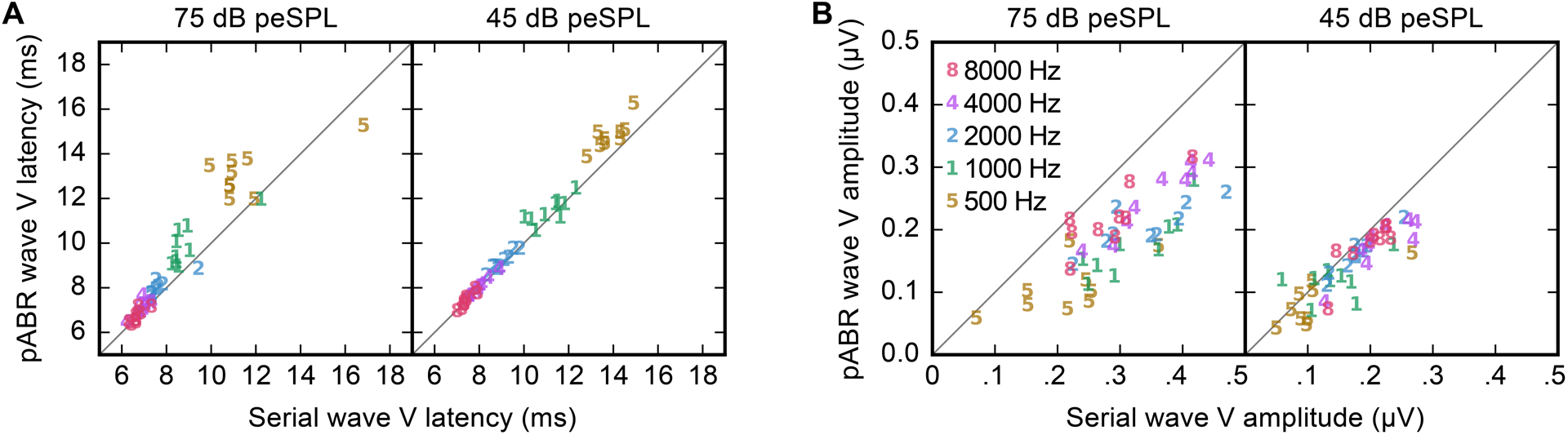
Comparison of pABR with serial wave V latency (**A**) and amplitude (**B**) for all subjects (N = 9) at each frequency for both stimulus levels. Quantities shown are for the right ear. Stimulus frequency indicated by marker number and color.

### Acquisition times are faster for pABR than serial measurement

Having compared waveform morphology and demonstrated that pABR provides canonical waveforms with only minor systematic differences in wave V amplitude and latency, we next compared the acquisition time of pABR to serial measurements.

First we characterized the time (in minutes) it took to reach a residual noise of 20 nV, which was calculated for each waveform as the square root of 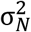. For pABR recording, the time for the responses across all frequencies to reach criterion was defined as the time taken by the slowest response to reach 20 nV (i.e., the maximum time across all responses). For serial measurement, the acquisition time was the sum of the times for each of the five frequencies’ responses to reach criterion, doubled to account for the other ear. We calculated the time to the 20 nV residual noise criterion for all subjects at both stimulus levels, leading to 18 estimates for each acquisition method, which are plotted as a histogram in Figure 8. The pABR reached 20 nV for all waveforms with a median time of 4.6 minutes (3.8–5.4 minutes interquartile range). Serial recordings, on the other hand, took substantially longer at 30.1 minutes (23.8–40.0 minutes). Dividing each serial time by each corresponding pABR time yielded a median speedup ratio of 6.0 (5.7–6.6 interquartile range), indicating a large advantage for the pABR.

**Figure 8.**
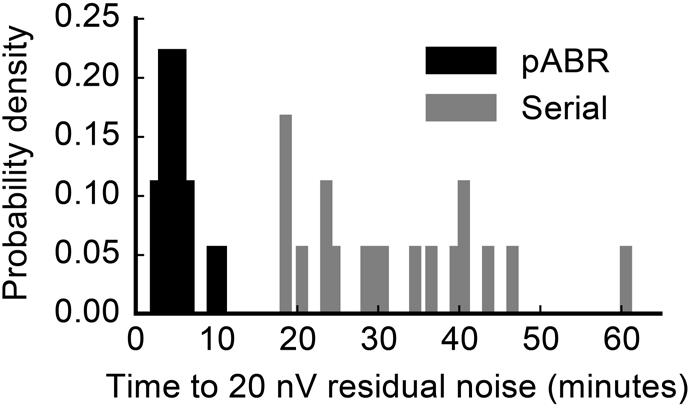
Comparison of time to reach 20 nV residual noise for all 10 waveforms between methods (pABR in black, serial in gray). Two intensities for 9 subjects are shown, leading to 18 data points for each acquisition method.

The residual noise numbers indicate that in situations where multiple waveforms are desired, recording them in parallel leads to lower noise levels much faster than recording them one at a time. If pABR yielded identical responses to serial measurement, then the speedup ratios for response acquisition would be higher. However, because the pABR leads to smaller responses in some situations (low frequencies at high intensities), the speedups were less pronounced, particularly at higher intensities.

Second, we further compared estimated acquisition times by calculating the time required for all waveforms of a given intensity to reach 0 dB SNR^1^. As with the residual noise estimates, the total acquisition time for pABR was the time it took for the last waveform to reach threshold, and the total time for serial acquisition was the sum of acquisition times for all waveforms (here approximated as the total for one ear, doubled). These times are given for all subjects in Table 1, and Figure 9A shows an example acquisition run modeled from one subject’s data for demonstration purposes. At 75 dB peSPL, the median acquisition time for pABR was 1.93 minutes (0.93–3.63 minutes interquartile range) and for serial acquisition was 1.45 minutes (0.94–3.45 minutes interquartile range). Parallel acquisition was faster for 5 of 9 subjects, with a median pABR speedup ratio of 1.45 (0.89–1.64). At 45 dB peSPL the acquisition time difference was pronounced: median acquisition time for pABR was 4.60 minutes (1.86–8.99 minutes) versus 7.81 minutes (5.95–9.92 minutes) for serial presentation. At this lower intensity, pABR was faster than serial recording for all 9 subjects, with a median pABR speedup ratio of 2.99 (1.12–3.92 interquartile range). A scatterplot comparing the pABR and serial acquisition times at 75 and 45 dB peSPL (filled and open circles, respectively) is shown in Figure 9B. Points below the unity line indicate a pABR advantage.

**Figure 9.**
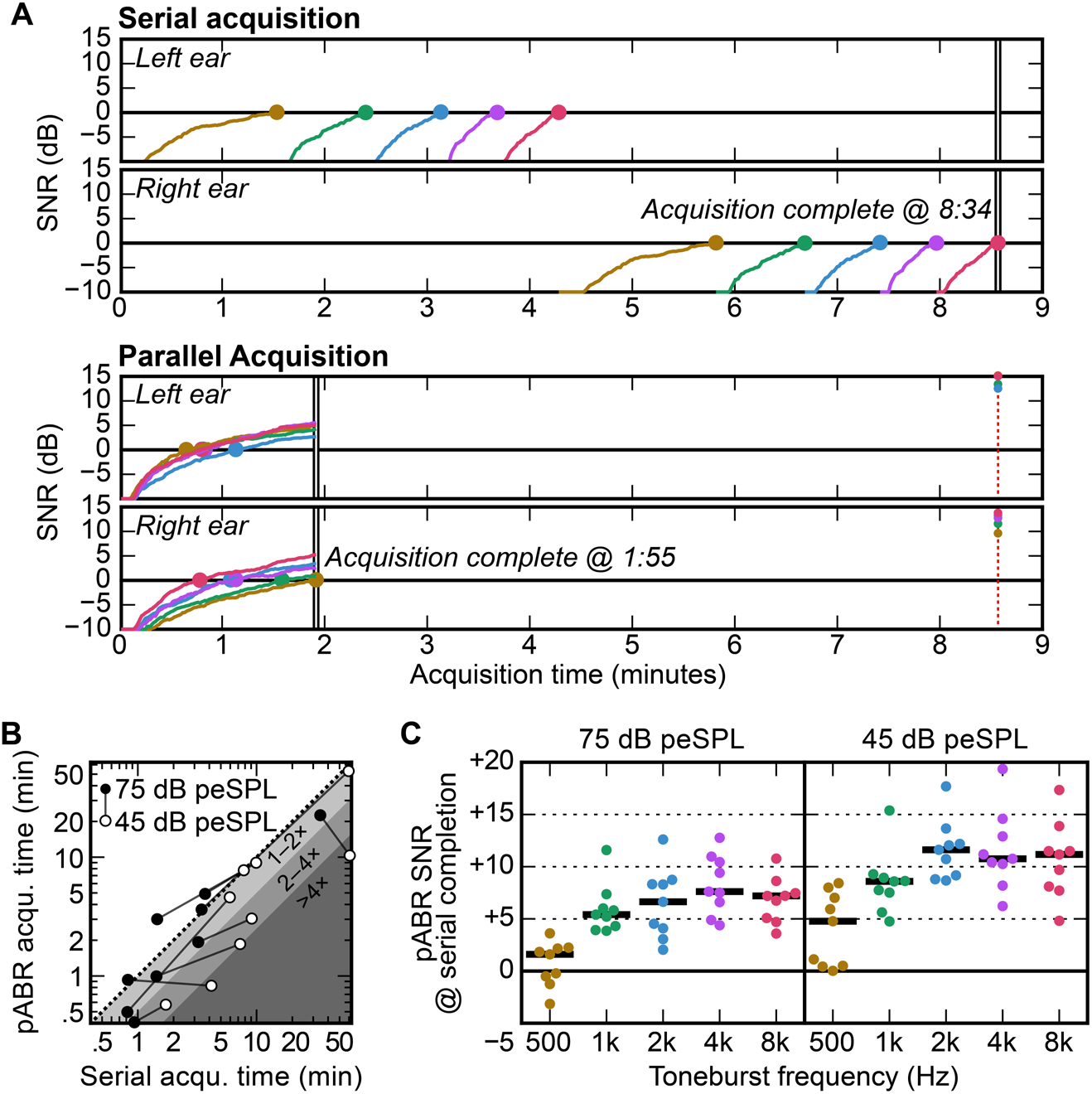
pABR shows faster acquisition and better SNR. (**A**) Real-time acquisition runs simulated from offline data for one subject at 45 dB peSPL. For serial ABR, unrecorded left ear runs were assumed to be equal to right ear. (**B**) Comparison recording time for 9 subjects. Points below dotted unity line are cases where pABR is faster. Shaded regions indicate speedup ratios of 1–2 (light gray), 2–4 (medium gray), and > 4 (dark gray). (**C**) SNR of pABR runs upon serial acquisition completion (subjects colored points, median black lines), corresponding for one subject to the points on the red dashed vertical line in **A**.

**Table 1.**
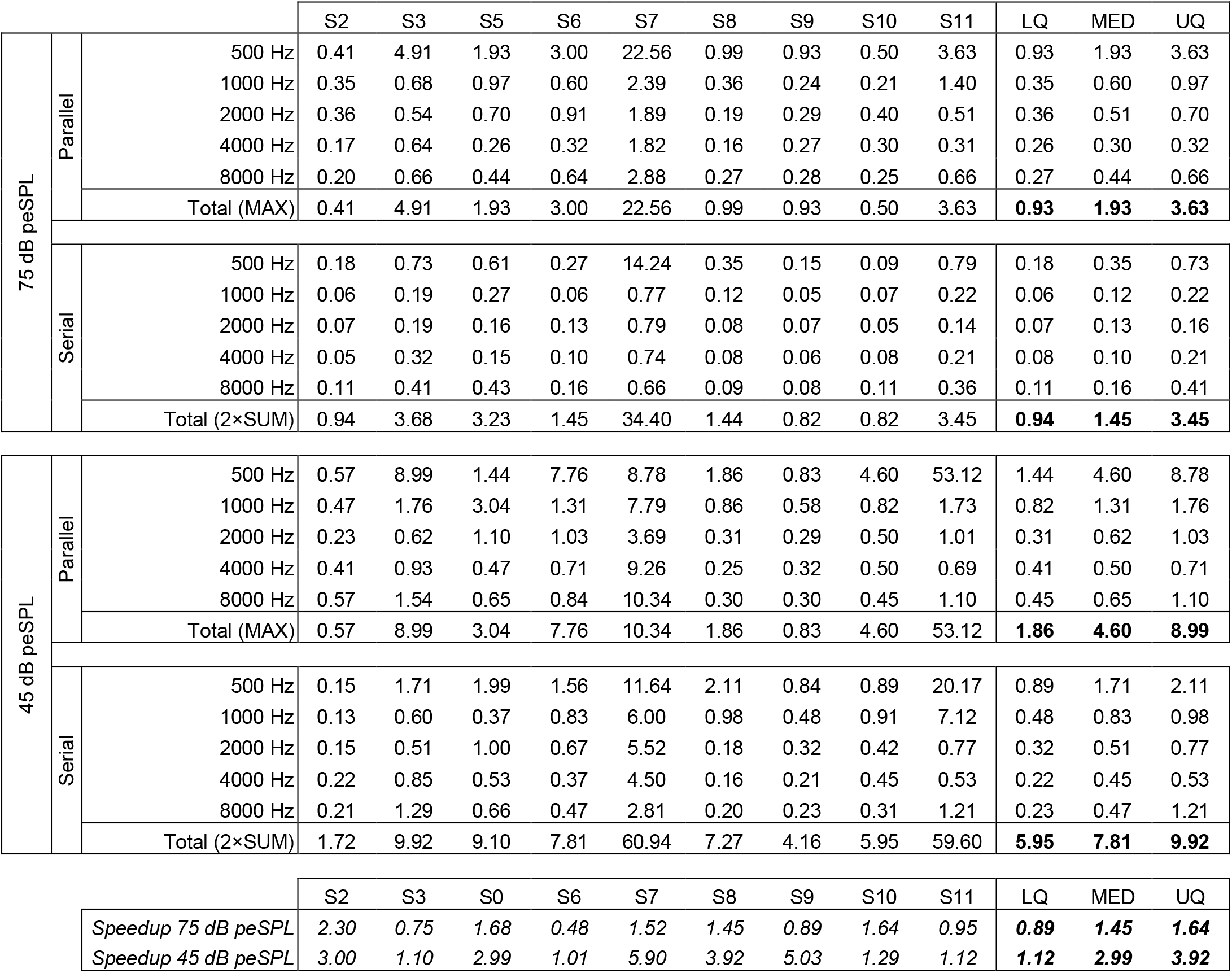
Time to 0 dB SNR (in minutes) for each subject as well as the median and quantiles. For each method at each stimulus level the time is shown for all frequencies, with the total (computed with the appropriate method) shown below. Shown at the bottom in italics are the speedups for both stimulus levels. These numbers are unitless ratios, rather than minutes, where higher numbers represent an advantage for the pABR over the serial ABR.

|  |  |  | S2 | S3 | S5 | S6 | S7 | S8 | S9 | S10 | S11 | LQ | MED | UQ |
| --- | --- | --- | --- | --- | --- | --- | --- | --- | --- | --- | --- | --- | --- | --- |
| 75 dB peSPL | Parallel | 500 Hz | 0.41 | 4.91 | 1.93 | 3.00 | 22.56 | 0.99 | 0.93 | 0.50 | 3.63 | 0.93 | 1.93 | 3.63 |
|  |  | 1000 Hz | 0.35 | 0.68 | 0.97 | 0.60 | 2.39 | 0.36 | 0.24 | 0.21 | 1.40 | 0.35 | 0.60 | 0.97 |
|  |  | 2000 Hz | 0.36 | 0.54 | 0.70 | 0.91 | 1.89 | 0.19 | 0.29 | 0.40 | 0.51 | 0.36 | 0.51 | 0.70 |
|  |  | 4000 Hz | 0.17 | 0.64 | 0.26 | 0.32 | 1.82 | 0.16 | 0.27 | 0.30 | 0.31 | 0.26 | 0.30 | 0.32 |
|  |  | 8000 Hz | 0.20 | 0.66 | 0.44 | 0.64 | 2.88 | 0.27 | 0.28 | 0.25 | 0.66 | 0.27 | 0.44 | 0.66 |
|  |  | Total (MAX) | 0.41 | 4.91 | 1.93 | 3.00 | 22.56 | 0.99 | 0.93 | 0.50 | 3.63 | **0.93** | **1.93** | **3.63** |
|  | Serial | 500 Hz | 0.18 | 0.73 | 0.61 | 0.27 | 14.24 | 0.35 | 0.15 | 0.09 | 0.79 | 0.18 | 0.35 | 0.73 |
|  |  | 1000 Hz | 0.06 | 0.19 | 0.27 | 0.06 | 0.77 | 0.12 | 0.05 | 0.07 | 0.22 | 0.06 | 0.12 | 0.22 |
|  |  | 2000 Hz | 0.07 | 0.19 | 0.16 | 0.13 | 0.79 | 0.08 | 0.07 | 0.05 | 0.14 | 0.07 | 0.13 | 0.16 |
|  |  | 4000 Hz | 0.05 | 0.32 | 0.15 | 0.10 | 0.74 | 0.08 | 0.06 | 0.08 | 0.21 | 0.08 | 0.10 | 0.21 |
|  |  | 8000 Hz | 0.11 | 0.41 | 0.43 | 0.16 | 0.66 | 0.09 | 0.08 | 0.11 | 0.36 | 0.11 | 0.16 | 0.41 |
|  |  | Total (2×SUM) | 0.94 | 3.68 | 3.23 | 1.45 | 34.40 | 1.44 | 0.82 | 0.82 | 3.45 | **0.94** | **1.45** | **3.45** |

| 45 dB peSPL | Parallel | 500 Hz | 0.57 | 8.99 | 1.44 | 7.76 | 8.78 | 1.86 | 0.83 | 4.60 | 53.12 | 1.44 | 4.60 | 8.78 |
| --- | --- | --- | --- | --- | --- | --- | --- | --- | --- | --- | --- | --- | --- | --- |
|  |  | 1000 Hz | 0.47 | 1.76 | 3.04 | 1.31 | 7.79 | 0.86 | 0.58 | 0.82 | 1.73 | 0.82 | 1.31 | 1.76 |
|  |  | 2000 Hz | 0.23 | 0.62 | 1.10 | 1.03 | 3.69 | 0.31 | 0.29 | 0.50 | 1.01 | 0.31 | 0.62 | 1.03 |
|  |  | 4000 Hz | 0.41 | 0.93 | 0.47 | 0.71 | 9.26 | 0.25 | 0.32 | 0.50 | 0.69 | 0.41 | 0.50 | 0.71 |
|  |  | 8000 Hz | 0.57 | 1.54 | 0.65 | 0.84 | 10.34 | 0.30 | 0.30 | 0.45 | 1.10 | 0.45 | 0.65 | 1.10 |
|  |  | Total (MAX) | 0.57 | 8.99 | 3.04 | 7.76 | 10.34 | 1.86 | 0.83 | 4.60 | 53.12 | **1.86** | **4.60** | **8.99** |
|  | Serial | 500 Hz | 0.15 | 1.71 | 1.99 | 1.56 | 11.64 | 2.11 | 0.84 | 0.89 | 20.17 | 0.89 | 1.71 | 2.11 |
|  |  | 1000 Hz | 0.13 | 0.60 | 0.37 | 0.83 | 6.00 | 0.98 | 0.48 | 0.91 | 7.12 | 0.48 | 0.83 | 0.98 |
|  |  | 2000 Hz | 0.15 | 0.51 | 1.00 | 0.67 | 5.52 | 0.18 | 0.32 | 0.42 | 0.77 | 0.32 | 0.51 | 0.77 |
|  |  | 4000 Hz | 0.22 | 0.85 | 0.53 | 0.37 | 4.50 | 0.16 | 0.21 | 0.45 | 0.53 | 0.22 | 0.45 | 0.53 |
|  |  | 8000 Hz | 0.21 | 1.29 | 0.66 | 0.47 | 2.81 | 0.20 | 0.23 | 0.31 | 1.21 | 0.23 | 0.47 | 1.21 |
|  |  | Total (2×SUM) | 1.72 | 9.92 | 9.10 | 7.81 | 60.94 | 7.27 | 4.16 | 5.95 | 59.60 | **5.95** | **7.81** | **9.92** |

**Table 1.** Time to 0 dB SNR (in minutes) for each subject as well as the median and quantiles.
|  |  |  | S2 | S3 | S0 | S6 | S7 | S8 | S9 | S10 | S11 | LQ | MED | UQ |
| --- | --- | --- | --- | --- | --- | --- | --- | --- | --- | --- | --- | --- | --- | --- |
| Speedup 75 dB peSPL |  |  | 2.30 | 0.75 | 1.68 | 0.48 | 1.52 | 1.45 | 0.89 | 1.64 | 0.95 | **0.89** | **1.45** | **1.64** |
| Speedup 45 dB peSPL |  |  | 3.00 | 1.10 | 2.99 | 1.01 | 5.90 | 3.92 | 5.03 | 1.29 | 1.12 | **1.12** | **2.99** | **3.92** |
| 45 dB peSPL | Parallel | 500 Hz | 0.57 | 8.99 | 1.44 | 7.76 | 8.78 | 1.86 | 0.83 | 4.60 | 53.12 | 1.44 | 4.60 | 8.78 |
|  |  | 1000 Hz | 0.47 | 1.76 | 3.04 | 1.31 | 7.79 | 0.86 | 0.58 | 0.82 | 1.73 | 0.82 | 1.31 | 1.76 |
|  |  | 2000 Hz | 0.23 | 0.62 | 1.10 | 1.03 | 3.69 | 0.31 | 0.29 | 0.50 | 1.01 | 0.31 | 0.62 | 1.03 |
|  |  | 4000 Hz | 0.41 | 0.93 | 0.47 | 0.71 | 9.26 | 0.25 | 0.32 | 0.50 | 0.69 | 0.41 | 0.50 | 0.71 |
|  |  | 8000 Hz | 0.57 | 1.54 | 0.65 | 0.84 | 10.34 | 0.30 | 0.30 | 0.45 | 1.10 | 0.45 | 0.65 | 1.10 |
|  |  | Total (MAX) | 0.57 | 8.99 | 3.04 | 7.76 | 10.34 | 1.86 | 0.83 | 4.60 | 53.12 | **1.86** | **4.60** | **8.99** |
|  | Serial | 500 Hz | 0.15 | 1.71 | 1.99 | 1.56 | 11.64 | 2.11 | 0.84 | 0.89 | 20.17 | 0.89 | 1.71 | 2.11 |
|  |  | 1000 Hz | 0.13 | 0.60 | 0.37 | 0.83 | 6.00 | 0.98 | 0.48 | 0.91 | 7.12 | 0.48 | 0.83 | 0.98 |
|  |  | 2000 Hz | 0.15 | 0.51 | 1.00 | 0.67 | 5.52 | 0.18 | 0.32 | 0.42 | 0.77 | 0.32 | 0.51 | 0.77 |
|  |  | 4000 Hz | 0.22 | 0.85 | 0.53 | 0.37 | 4.50 | 0.16 | 0.21 | 0.45 | 0.53 | 0.22 | 0.45 | 0.53 |
|  |  | 8000 Hz | 0.21 | 1.29 | 0.66 | 0.47 | 2.81 | 0.20 | 0.23 | 0.31 | 1.21 | 0.23 | 0.47 | 1.21 |
|  |  | Total (2×SUM) | 1.72 | 9.92 | 9.10 | 7.81 | 60.94 | 7.27 | 4.16 | 5.95 | 59.60 | **5.95** | **7.81** | **9.92** |

**Table 1.** Time to 0 dB SNR (in minutes) for each subject as well as the median and quantiles.
|  |  |  | S2 | S3 | S0 | S6 | S7 | S8 | S9 | S10 | S11 | LQ | MED | UQ |
| --- | --- | --- | --- | --- | --- | --- | --- | --- | --- | --- | --- | --- | --- | --- |
| Speedup 75 dB peSPL |  |  | 2.30 | 0.75 | 1.68 | 0.48 | 1.52 | 1.45 | 0.89 | 1.64 | 0.95 | **0.89** | **1.45** | **1.64** |
| Speedup 45 dB peSPL |  |  | 3.00 | 1.10 | 2.99 | 1.01 | 5.90 | 3.92 | 5.03 | 1.29 | 1.12 | **1.12** | **2.99** | **3.92** |

Even in the case where a pABR and corresponding serial acquisition take the same amount of time, there is a secondary SNR advantage for pABR which comes from the criterion that ends each run. For the pABR, data continues to accrue for all waveforms while waiting for the last response to reach criterion. Therefore, at the end of the run all but the slowest waveform will have an SNR better than the stopping criterion. In contrast, for serial acquisition, at the end of the run all waveforms will have just reached criterion SNR. This difference can be seen in Figure 9A, where at the end of the parallel run (double black line at time 1:55) all but the 500 Hz right ear response are better than 0 dB SNR. This pABR SNR benefit can be quantified by examining the SNR of the pABR waveforms at the time point when the corresponding serial run completed (red dashed line at time 8:34). These SNR benefits are plotted in Figure 9C for all frequencies at both intensities. At 75 dB peSPL, the median SNR benefits for 500 through 8000 Hz are 1.6, 5.4, 6.6, 7.6, 7.2 dB. For lower intensity of 45 dB peSPL, the benefits are even greater: 4.8, 8.6, 11.6, 10.7, 11.2 dB from 500 to 8000 Hz. These improvements potentially allow much better assessment of waveform morphology, such as the presence and size of wave I.

## DISCUSSION

Here we describe the pABR, a new method for recording the frequency-specific ABR to multiple simultaneous stimulus trains at several octave frequencies in both ears. The pABR yields waveforms with canonical response components, namely wave V, albeit at slightly different latencies and amplitudes at higher intensities. The principal advantage of the pABR is that low noise levels are achieved in drastically shorter times, which leads to faster acquisition times. Faster response acquisition will yield shorter clinic visits, or visits of the same length that yield much better estimates of the hearing thresholds on which crucial clinical decisions are based. Furthermore, octave frequencies from 500 to 8000 Hz can be obtained in comparable or shorter lengths of time, which provides a more comprehensive assessment of hearing function than typically achieved in current clinical practice. At best, 500–4000 Hz thresholds are currently achieved but more often only 500, 2000 and maybe 4000 Hz are obtained (American Academy of Audiology, 2012; BC Early Hearing Program, 2012; Hyde, 2008). The most obvious question about the pABR—how *much* faster it is—is also the most difficult to answer because it depends on a multitude of factors. We discuss several of these factors below.

The estimate of acquisition time we used here—time to criterion SNR—was objective but most useful for relative comparisons between the methods rather than absolute estimates of acquisition time. First, the choice of SNR has a large effect on the time in minutes (using +3 dB instead of 0 dB would have doubled all the times, for instance), so the times reported here should be considered within the context of our chosen criterion. However, a change in criterion would not affect the speedup ratios. These ratios indicate that the pABR can yield 10 good waveforms about 3 times faster than the serial ABR in at least half the cases (Table 1). This means more information could be collected in an appointment, or the same amount of information could be collected quicker. For some subjects, acquisition of 10 waveforms occurred quickly, with times as low as less than half a minute (Table 1). For the pABR at a level closer to threshold (i.e., 45 dB peSPL), 5 of 9 subjects achieved good waveforms within 5 minutes, compared to only 2 subjects with serial presentation. As would happen in the clinic, there were some subjects that had noisier responses and took substantially longer to acquire 10 waveforms with both parallel and serial presentation, such as two subjects who achieved waveforms in estimated times of about 53 (pABR) and 61 (serial) minutes. The pABR is subject to the effects of noisy testing situations, just as the serial ABR. However, these time estimates may also be conservative given the automatic calculation of SNR. Importantly, audiologists are highly trained at recognizing response components. For example, in many cases while analyzing our data we could see a clear 500 Hz response when the SNR was still below our 0 dB SNR criterion. For those subjects who had estimated times to 0 dB SNR greater than 10 minutes, a trained audiologist would likely detect the presence or absence of a waveform earlier and make decisions about moving on to another level. Testing with trained clinicians interpreting waveforms as they are acquired in real time will give more meaningful time estimates in minutes.

The pABR offers advantages that will make clinicians’ decisions about response presence more accurate and easier to make. First, viewing the response to a specific frequency in context of the other frequencies being simultaneously acquired allows the clinician to make a better, holistic assessment of its presence/absence than viewing the same waveform in isolation. Second, extending the analysis window (made possible by the random stimulus timing) can show later response components, such as the middle latency response (MLR), which when present can further eliminate uncertainty whether a response is present or absent. Our focus was on the ABR, but extending the signal beyond 10 ms to include the MLR will improve SNR estimates and may also further decrease acquisition times based on time to 0 dB SNR. Including the MLR may have shortened the long acquisition times estimated for the particularly noisy subjects discussed above (see Table 1). The pre-stimulus period can also be extended, giving a better impression of the noise. These advantages are highlighted in Figure 10. In panel A, the 500 Hz response is shown on its own. A response may be present, but its amplitude is only slightly greater than that of the noise. In panel B, the same response is shown along with the other simultaneously recorded frequencies for that ear, making the 500 Hz response easier to see. In panel C, the extended analysis window provides a clearer pre-stimulus baseline and middle latency components at ~35 ms that make the presence of a 500 Hz response more certain. The use of latencies beyond the typical ABR window will be an important subject of future investigation.

**Figure 10.**
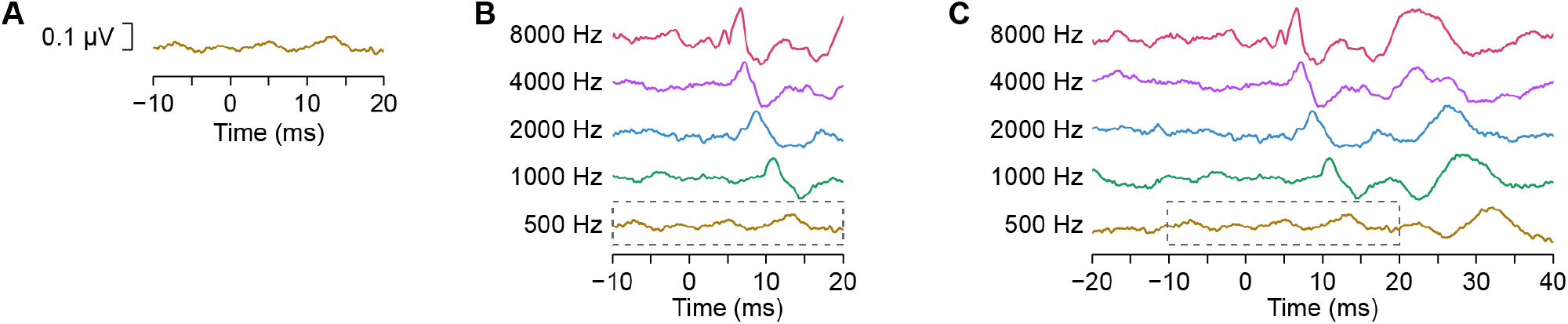
Improved visual response detection. (**A**) 500 Hz response waveform alone. (**B**) Same response with other frequencies simultaneously acquired. Dotted gray box surrounds the waveform from **A**. (**C**) 500 Hz response with other frequencies present and analysis window extended 10 ms to the left and 20 ms to the right. Gray box as in **B**.

**Figure 11.**
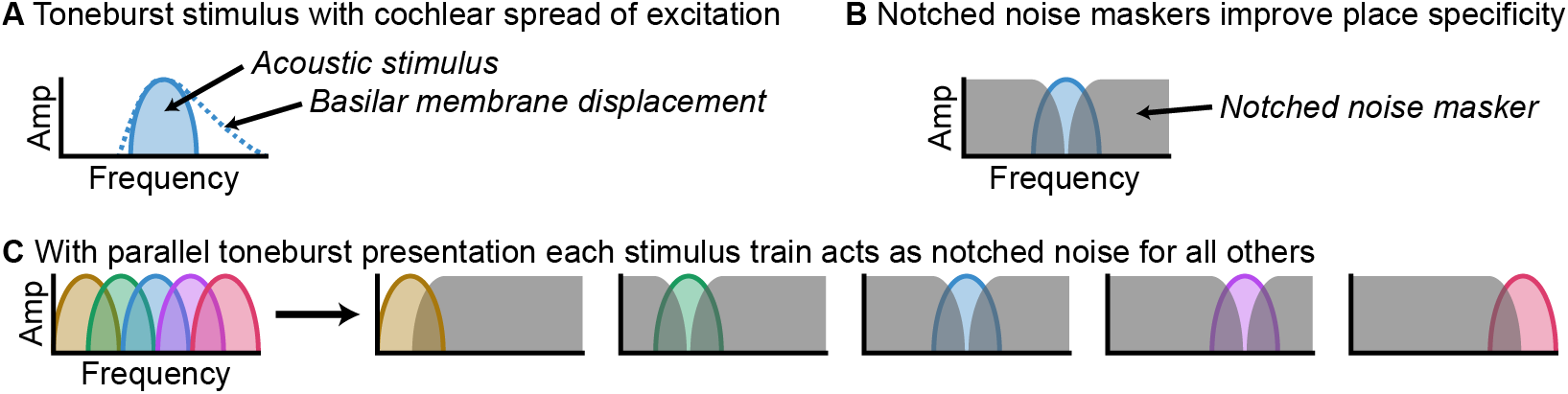
Parallel stimulation allows toneburst trains to also function as notched noise. (**A**) Cartoon representation of a single toneburst stimulus in the frequency domain (filled area), and the pattern of excitation it evokes in the cochlea (dashed line), showing spread of excitation towards the basal end. (**B**) Notched noise can be used to mask the off-frequency excitation, yielding a more place-specific response. (**C**) In pABR, each frequency band is masked by the others, which summed have a similar effect to a notched noise masker.

The hearing thresholds of the people being tested will also have a large effect on the overall measurement time. In the present study all subjects had normal hearing thresholds, and 500 Hz was the most difficult response to acquire. This is not surprising, and even during current diagnostic exams normal hearing for 500 Hz is determined using a higher level than the other frequencies (American Academy of Audiology, 2012; BC Early Hearing Program, 2012; Hyde, 2008). However, high frequency sloping loss is the most common configuration (e.g., Pittman & Stelmachowicz, 2003). Thus, while the pABR’s speed advantage was limited by the low frequency acquisition time here, this may not be true in most cases where a hearing loss is present. At higher levels necessary to determine high frequency thresholds, the level for 500 Hz would be suprathreshold and generate responses with larger SNRs quicker than at a level near threshold (e.g., time to 0 dB SNR for 75 versus 45 dB peSPL in Table 1, Figure 9B). Consequently, the actual acquisition time could be further reduced relative to traditional methods. Additionally, because the pABR reaches low residual noise levels faster than traditional methods, the pABR may allow clinicians to more quickly determine “no response” when none is present.

There are thus several factors that limit our ability to fully predict the absolute speed gains the pABR will provide in the clinic. Even non-measurement times between runs will be reduced because the clinician need only select the next intensity to test, rather than choosing a specific intensity-frequency-ear combination as the next step of the threshold search. We show here that the pABR is faster than traditional methods and offers a number of factors that may further improve the speedup. The next step to quantifying the full advantages for clinical use will involve testing the pABR in an actual clinical setting with the patients of interest—namely people (adults, infants, and children) with a wide range of hearing loss.

The pABR is not the only objective audiometric tool that allows simultaneous threshold estimation at multiple frequencies—this is also accomplished by the multiple ASSR. As such, the ASSR warrants comparison with the pABR. The ASSR is an evoked response that is phase-locked to a periodic stimulus and can also be measured with most ABR hardware. In clinical settings, the stimulus is typically a tonal carrier at the test frequency (e.g. 500 Hz) whose amplitude is modulated to create the steady-state response. Modulation frequencies in the 80–100 Hz range are used to avoid contributions from cortical generators which are affected by subject state (Korczak, Smart, Delgado, Strobel, & Bradford, 2012). As with the toneburst ABR, correlations between ASSR and behavioral thresholds reach around 0.9 when a large range is considered (Luts et al., 2006). More than one frequency and ear can be tested at a time by “tagging” them with different modulator frequencies. Rather than waveforms, however, the ASSR assessment is based on a scalar measure of its phase-locking to the modulator (and its harmonics), expressed as a single summary quantity. In contrast, the pABR provides full response waveforms. This carries a number of advantages: 1) it allows inference beyond the presence or absence of a response, such as the investigation of auditory neuropathy and site-of-lesion testing, 2) it allows the separation of brainstem and cortical responses by their latencies, letting the clinician use middle latency cortical responses if present, and 3) it will require less training because it draws on clinicians’ existing expertise in interpreting ABR waveforms.

Because the pABR tests multiple frequencies at once, the potential for interactions between stimuli in the cochlea must be considered. Even though highly frequency-specific stimuli can be generated, they may not elicit place-specific displacements along the basilar membrane when presented without masking (as is typical). High intensity stimuli elicit broader excitation patterns (Robles & Ruggero, 2001) and excitation asymmetrically spreads towards the base of the cochlea. Therefore, responses to low-frequency stimuli include greater contributions from other parts of the cochlea with higher best frequencies. However, the pABR has the potential to provide better place-specific responses because each of the frequency bands could act as masking noise for all the others, as depicted in Figure 8. Essentially, the pABR could act akin to recording a series of masked ABRs but in one run. Evidence that may support this place-specific hypothesis comes from the prolonged latencies for the pABR relative the serially recorded responses, especially for the lower frequencies at higher intensities (Figures 6 and 7). Because spread of excitation is greater at higher intensities, we would expect to see the biggest differences between pABR and serial ABR at higher levels. Indeed, we found that at the lower level of 45 dB peSPL, there was no difference between the two methods in wave V amplitude or latency, indicating minimal interference. However, at the higher stimulus level of 75 dB peSPL, wave V amplitude was reduced and wave V latency longer for pABR than serial ABR for the lower frequencies. These differences for lower frequencies are consistent with basal spread of activation contributing to responses in the traditional ABR but being masked under the pABR. Thus, at lower stimulus levels, where acquisition generally takes longer and speedups are the most needed, interactions between bands of the cochlea do not seem to be an issue. At higher levels interactions appear to be present, likely leading to more modest speedups, but potentially in exchange for (or because of) improved place specificity.

In summary, the pABR is a viable method for recording canonical ABR waveforms at a fraction of the time of traditional serial methods, particularly for lower intensity stimuli. Consequently, the pABR has great potential for facilitating quick and accurate hearing threshold estimation that is important for timely diagnosis and treatment of hearing loss. Furthermore, the advantages of extended analysis windows afforded by randomized timing allows better noise estimates and inclusion of additional peaks such as the MLR, which will improve SNR estimates. Finally, our results suggest that the masking provided by simultaneously presented tonebursts might mitigate spread of activation at higher intensities, with potential improvements in place specificity. Future studies will focus on investigating optimal parameters for the pABR to estimate thresholds, modeling place specificity of the pABR, and assessing the utility of the pABR for estimating thresholds for various configurations of hearing loss with patients in the clinic.

## ACKNOWLEDGEMENTS

The authors wish to thank Sara Fiscella and Madeline Cappelloni for assistance with data collection, Veronica Valencerina and Kevin Paskiet for assistance with piloting, and Mark Orlando for many helpful discussions.

## FUNDING

This work was supported by National Institute for Deafness and Other Communication Disorders grant [R00DC014288] awarded to RKM.

## DATA AVAILABILITY

Data will be made available upon reasonable request to the corresponding author.

## Footnotes

1 The choice of 0 dB as the SNR threshold was arbitrary and based on visual assessment of when waveforms looked “good.” Changing this threshold would have changed the acquisition times. This change, however, would be multiplicative, such that the speedup ratios–our measure of how much faster the pABR is–would be unaffected.

## Notes

#### Summary of Updates

Slightly updated methods.

